# Antibiotic-induced dysbiosis does not potentiate the effect of lipopolysaccharide insult in male Wistar rats

**DOI:** 10.1101/2022.12.29.522253

**Authors:** Hana Tejkalová, Lea Jakob, Simona Kvasnová, Jan Klaschka, Hana Sechovcová, Jakub Mrázek, Tomáš Páleníček, Kateřina Olša Fliegerová

## Abstract

The present study investigated whether neonatal exposure to the proinflammatory endotoxin lipopolysaccharide (LPS) followed by antibiotic (ATB)-induced dysbiosis in early adulthood could induce schizophrenia-like behavioral changes in adult male rats. The combination of these two stressors resulted in decreased weight gain, but no significant behavioral abnormalities were observed. LPS treatment resulted in adult rats hypoactivity and induced anxiety-like behavior in the social recognition paradigm, but these behavioral changes were not exacerbated by ATB-induced gut dysbiosis. ATB treatment seriously disrupted the gut bacterial community, but dysbiosis did not affect locomotor activity, social recognition, and acoustic reactivity in adult rats. Fecal bacterial community analyzes showed no differences between the LPS challenge exposed/unexposed rats, while the effect of ATB administration was decisive regardless of prior LPS exposure. ATB treatment resulted in significantly decreased bacterial diversity, suppression of Clostridiales and Bacteroidales, and increases in Lactobacillales, Enterobacteriales, and Burkholderiales. The persistent effect of LPS on some aspects of behavior suggests a long-term effect of early toxin exposure that was not observed in ATB-treated animals. However, an anti-inflammatory protective effect of ATB cannot be assumed because of the increased abundance of pro-inflammatory, potentially pathogenic bacteria (*Proteus, Suttrella*) and the elimination of the bacterial families *Ruminococcaceae* and *Lachnospiraceae*, which are generally considered beneficial for gut health.

## 1. Introduction

The current state of the art suggests an important communication between the residing intestinal microorganisms, brain development and function, and also behavior associated with psychiatric disorders [1–6]. The concept known as the microbiota-gut-brain axis is now widely accepted [7–13], and in neuroscience, the number of studies on the interactions between the brain, gastrointestinal tract, gut bacteria, and the bidirectional relationship between these systems is steadily increasing. The rodent model of psychiatric disorders is a valuable source of information that can greatly expand our understanding of the role of the gut microbiota in the etiopathology of mental diseases and elucidate the effects of the microbiome on behavior and cognitive function [14,15]. Exposure of rodents to inflammatory agents, such as the proinflammatory endotoxin lipopolysaccharide (LPS), during early development is one of the animal models used to study neurodevelopmental disorders such as autism and schizophrenia [16,17]. LPS is part of the outer membrane of Gram-negative bacteria and is known to induce a strong immune response, resulting in increased levels of proinflammatory mediators (e.g., IL -1β, TNF-α) that affect the brain [18] and induce sickness behavior [19].

Increased anxiety-like and depressive-like behavior after LPS administration have been reported in mice [20–23] and rats [24–29]. In these works, a decrease in locomotor activity, food intake, social interaction, and/or exploration for novel objects was observed in rodents several hours after the endotoxin insult. Studies on the long-term effects of LPS treatment are not as consistent, showing increased anxiety-like behavior [30], compulsive responses [31], and impaired social interaction [32], while no impact on locomotor activity were observed in rats in rats [17,31] and no significant changes in anxiety- and depression-like behaviors were observed in adult mice [33]. On the other hand, studies mostly agree that LPS leads to disruption of sensorimotor gating [17,34–37] expressed as the prepulse inhibition (PPI) deficit, a proposed biomarker of schizophrenia-like behavior [38–40].

Although neonatal LPS immune activation is considered a crucial factor in vulnerability to neuropsychopathological disorders, Bilbo et al. [41] observed in rat model, that some imparment effects may not occur until a secondary stressor is introduced. In particular, complex neurodevelopmental disorders such as schizophrenia or autism may require both exposures, i.e., a specific infection at a specific time in early development combined later with other environmental hit [42]. The type of stressor is important and significantly affects outcome [43]. The second hit can include secondary immune challenge [41], different kind of stress [30,44,45] or gut dysbiosis induced by antibiotics [11,46].

Antimicrobial treatment can be applied to induce microbiome perturbation under controlled conditions and can therefore be used in animals as a methodological tool to evaluate the effects of the gut microbiota on behavior [14,47]. In mice and rats, antibiotic administration in early life [48–50], adolescence [51] and adulthood [52–54] has been well studied and shown to alter many factors, including hormone levels, gene expression, anxiety-related responses, exploratory and social behaviors, and cognitive functions. On the other hand, only few papers have investigated the effect of LPS on the microbiome. This bacterial endotoxin is a potent activator of innate immune responses has been well studied and was shown to be able to changes many factors, including hormonal levels, gene expression, anxiety-related reaction, exploratory and social behavior, and cognitive functions. On the other hand, only few works investigated the effect of LPS on the microbiome. This bacterial endotoxin is powerful activator of innate immune responses [55], and LPS-induced cytokines are not derived only from innate or adaptive immune cells but also from intestinal epithelial cells and thus may be important factors in intestinal inflammation [56]. Gut microorganisms actively participate in the host inflammatory response by cytokine-microbiota or microbiota-cytokine modulation interactions [57]. This bidirectional relationship shapes the structure of the gut microbiota and changes in microbial composition can inhibit or stimulate inflammatory pathways [58].

However, the number of studies that have investigated the impact of LPS on the microbiome in laboratory animals is still limited and results vary between different rodent species. Significant effects on gut bacteria have been reported in mice [59–61], but no effect of toxin treatment on fecal bacterial community composition has been found in rats [62] or hamsters [46]. It is clear that the different rodent models are not comparable and that further research is needed to evaluate the consequences of an early inflammatory event on the gut microbiome.

Animal models are undoubtedly useful tools for studying the onset of neurodevelopmental disorders, explaining the origin of psychiatric abnormalities, creating model examples of pathological symptoms, or testing the effects of drugs that are not feasible in humans. Animal models are also critical for establishing cause-and-effect relationships between disease, behavior, and other factors, such as the microbiome [63]. The rodent model of schizophrenia is often based on the “two hit hypothesis”‘ formulated as early as 1999 by Bayer et al. [64]. This concept proposed that the development of the disease process requires a genetic predisposition (first hit) combined with the second hit(s) early or later in life, which may include multiple environmental factors such as viral infection(s), birth complications, or social stressors. This idea was reflected in animal models and led to a spate of rodent models of schizophrenia using several different first/second insults [42]. Prenatal or early life LPS challenge is often used as the first hit, but the types of second hit vary from study to study. Intestinal dysbiosis induced by antibiotic treatment is not commonly used, but this type of second insult is worth investigating because there is growing evidence that the mammalian digestive tract microbiome affects host physiology, modulates immunity, development, and nutrient uptake [65,66], and that its disruption affects not only physical but also mental health [8] due to the dysregulation of the gut-microbiome-brain axis [3,5,11,67–70].

The aim of the present study was to evaluate whether two stressors, the first applied in early life and the second in early adulthood, could induce schizophrenia-like behavioral changes in adulthood in a rodent model. Male rats were exposed to a combination of neonatal LPS challenge and antibiotic treatment in early adulthood to examine effects on locomotor activity, juvenile conspecific rat recognition, and sensorimotor gating. Behavioral tests were selected because they are known to be both relevant to symptoms of schizophrenia [71] and affected by changes in gut microbiota composition induced by antibiotic treatment [72]. As a secondary outcome, we determined the impact of both insults on the gut microbiome assessed by high-throughput sequencing (HTS) of 16S rRNA fragments of fecal bacteria. We hypothesize that the neonatal subchronic LPS insult has long-term effects on behavior and gut bacterial composition, but it is not clear whether antibiotic treatment in early adulthood can amplify or attenuate this presumed effect and induce schizophrenia-like behavior in rats.

## 2. Materials and Methods

### 2.1. Ethical Statement

The animal experiment was conducted in the facility of the National Institute of Mental Health (Klecany, Czech Republic). Animal handling, treatment, and clinical examinations were performed in accordance with instructions of the National Committee for the Care and Use of Laboratory Animals and were approved by the Local Animal Care Committee (MHCR No. 35/2017, 17668/2017-3OVZ, 13.4.2017). All procedures were performed in concordance with the European legislation Directive 2010/63/EU on the protection of animals used for scientific purposes and the Czech legislation Act on the Protection of Animals from Cruelty no. 246/1992.

### 2.2. Animals, Treatment and Study Design

One hundred fifty-three male Wistar/Hann rats (Velaz Ltd., Czech Republic) used in the experiments were housed in standard 48×21×21-cm polysulfone cages (Tecniplast, Buguggiate, Italy) in an air-conditioned room (22±2 °C) with a standard 12 h light/12 h dark cycle (lights on from 06:00-18:00). Rat litters with their mothers were obtained from the supplier on postnatal day 2 (PD 2). On PD 5 rats were permanently identified using finger marks, weighed, and the animals of each litter were divided into two cohorts. The first group was treated with lipopolysaccharide (LPS), an endotoxin from *E. coli*, serotype O26:B6 (L-8274, Sigma-Aldrich, St. Louis, MO, USA). LPS was dissolved in 0.9 % NaCl and administered intraperitoneally (i.p.) at a dose 2 mg/day/kg body weight (b.w.) to neonatal rats for 5 consecutive days (PD 5-9). A vehicle control containing 0.9 % NaCl was injected on the same days in the second group of animals that formed the control cohort (CTL). All pups were weaned on PD 28 and kept in smaller groups (3-4 animals) according to litter and treatment. Rats were fed normal chow diet (feed mixture ST-1, Velaz, Prague, Czech Republic) and were given tap water to drink ad libitum. From PD 50 to the end of the experiment, male rats were kept in pairs or a maximum of three animals per cage depending on neonatal pretreatment and adult treatments.

The endotoxin-treated group (LPS, n=79) and the saline-treated control group (CTL, n=74) were further randomly divided into LPS I (n=35) and CTL I (n=35) subgroups, which were transformed into two water-treated control groups LPS/W (n=35) and CTL/W (n=35). The other two subgroups LPS II (n=44) and CTL II (n=39) were transformed into antibiotic-treated (ATB) groups LPS/ATB (n=44) and CTL/ATB (n=39). The early adult rats in the antibiotic groups received the ATB mixture for 10 consecutive days in sterile drinking water (PD 60-69), whereas the rats in the control groups received the same amount of sterilized tap water (W). The antibiotic cocktail (50 ml/animal/day) consisted of metronidazole (500 mg/L; B. Braun, Melsungen, Germany), vancomycin (250 mg/L; Mylan S.A.S, Saint Priest), and colistine (15,000 U/mg; Sigma-Aldrich) to target broad spectrum of both Gram-positive and Gram-negative bacteria [73–75]. Drinking water in all cages was lightly sweetened with 10 g/L sucrose (Sigma-Aldrich) to suppress the bitter taste of the antibiotics. Drinking water was replaced daily, and water consumption in each cage was recorded. Animal body weights were recorded daily at PD 5-9 and subsequently at PD 16, PD 21, PD 28, PD 58, PD 60, and PD 70. Behavioral tests were performed as pretests before ATB or water treatment (PD 58-60) and as tests at the end of ATB or water treatment (PD 68-70). Feces were collected at the beginning (PD 60) and at the end of ATB or water treatment (PD 70). The schematic of the study design is shown in Figure 1.

**Figure 1.**
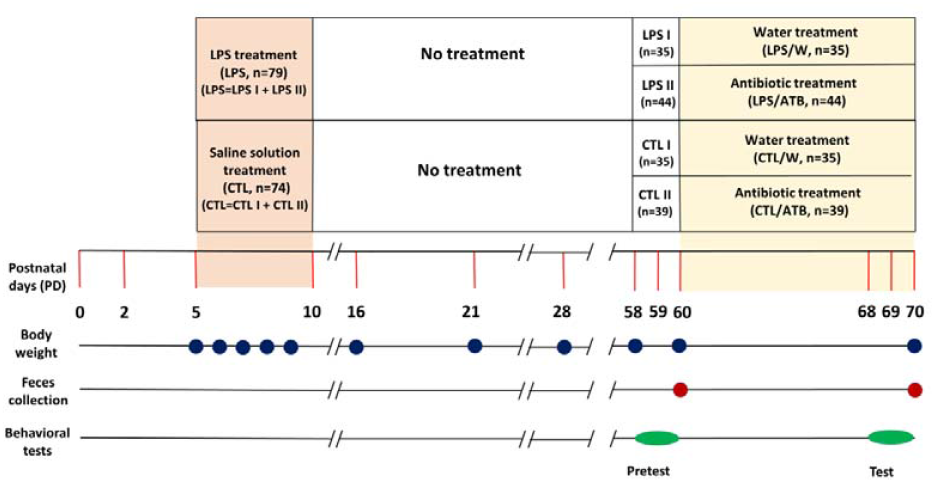
The study design diagram showing the experimental rat groups and time schedule of treatments (PD), body weight determinations (blue circles), fecal sampling (red circles), and behavioral pretests and tests (green ovals).

### 2.3. Behavioral Testing

Rats were tested in two blocks: Pretests (PD 58-60) and Tests (PD 68-70). The pretests were performed before the second stressor treatment to examine the effects of LPS toxin on the behavior of the early adult rats (CTL I vs LPS I, CTL II vs LPS II). The tests were performed at the end of ATB (or water) treatment to examine the long-term consequences of neonatal LPS insult alone (CTL/W vs LPS/W), of early adult ATB treatment alone (CTL/W vs CTL/ATB), and in combination with neonatal LPS hit (LPS/W vs LPS/ATB).

#### 2.3.1. Open Field Test

The Open Field Test (OFT) was used to assess locomotor and spontaneous exploratory activity. Locomotor activity was measured with a video camera connected to an Ethovision video tracking system (EthoVision Colour Pro-Version 3.1, Noldus Information Technology, Wageningen, The Netherlands). Rats were placed individually in the centre of the dimly lit arena (80 × 80 cm with black walls 30 cm high), which was located in a soundproof room, and locomotor activity was assessed as length of trajectory in cm over the 30-min series of trials (pretest and test). The arenas were cleaned between each trial.

#### 2.3.2. Social Recognition

The social recognition abilities of rats were assessed using the modified juvenile recognition procedure described by Richter et al [76]. Briefly, male rats tested were individually caged for 24 h prior to confrontation with a juvenile conspecific rat (22-26 days old) that represented a social stimulus. Immature rats were housed in small cages for 3 hours before being transferred to the home cage of the tested adult. Social behavior was assessed during a 5-min interaction period. Assessment was based on the adult’s exploration of the young rat’s head, body, and anogenital area, as well as forepaw manipulation and touching. After 30 minutes, the exposure of a young animal was repeated to determine whether the second exposure reflects juvenile-related memory. Testing was performed as a pretest before ATB administration and as a test at the end of ATB administration. The exploratory behavior of the adult towards the young animal was recorded and analyzed using the software ACTIVITIES [77]. The program generated pulses (at intervals of 0.1 s) and calculated the number of occurrences, total duration (i.e., total time spent performing the activity), and latency (i.e., time from onset of observation to first occurrence) of selected behavioral elements (recorded by key codes). In addition, rearing, a nonsocial parameter indicating in these circumstances the anxiety-like behavior was also documented. Experiments were scored by an experimenter who was blinded to the treatment groups.

#### 2.3.3. Acoustic Startle Response and Prepulse Inhibition

Acoustic Startle Response (ASR) and Prepulse Inhibition (PPI) tests were performed using the SR-LAB startle response system (San Diego Instruments, USA), as described previously [78]. Briefly, rats underwent a 5-min adaptation period in a startle chamber with continuous white noise background (70 dB). Then, 50 trials were presented in a pseudorandom order: 10 without stimulus for 30 ms, 30 prepulse + pulse inhibition with prepulse intensities of 3 (PP3), 5 (PP5), and 10 (PP10) dB[A] above background for 30 ms, and 10 startle pulse alone with intensity 120 dB (P120) for 30 ms. The interval between the prepulse and the startle pulse was fixed at 70 ms. The inter-trial period ranged from 10 to 20 s, and the entire session lasted 17 min. All animal groups were tested in a mixed order, always with litter-matched controls. ASR was assessed as the amplitude of the motor reaction expressed in mV. Prepulse inhibition was expressed as % PPI and calculated using the formula [100 - (100 × startle amplitude at PP)/P120], where PP is the average response to specific prepulse + pulse intensity trials and P120 is the startle alone. The total PP (PPx) was expressed as the mean of all different prepulses + pulses (PP3 + PP5 + PP10). Data from animals whose average response to the startle trials was below 10 mV (i.e., nonresponders) were excluded from the statistical calculations.

#### 2.3.4. Statistical Analysis

Behavioral tests and body weights were analyzed using BMDP and SigmaStat statistical software. All data, with some exceptions, were analyzed using a two-way between-subjects ANOVA with neonatal pretreatment as one factor and antibiotic treatment as the second factor. The exceptions are as follows. The pretest and test ASR and PPI values were compared using a three-way repeated-measures ANOVA model with two between-subjects factors of pretreatment (LPS vs. CTL), treatment (ATB vs. W), and one within-subjects factor of experimental phase (pretest vs test). The habituation curves resulting from the open field test were analyzed using a four-way repeated-measures ANOVA with two between-subjects factors of pretreatment and treatment, and two within-subjects factors of experimental phase, and time (6 levels, i.e., 5-minute segments). Finally, body weight analyses up to PD 60 were performed using a one-way between-subjects ANOVA with pretreatment as the only factor. Bonferroni-corrected Student t-tests were used as post-hoc tests. Test results were considered significant when p < 0.05. All data were expressed as means ± S.E.M.

### 2.4. Microbiome Analysis

#### 2.4.1. Sample Collection

Fresh fecal pellets were obtained aseptically by direct defecation of rats into a sterile 120-ml container (VWR, Radnor, PA, USA). Samples were collected from the 153 rats at PD 60, the pretest phase (baseline), and PD 70, the test phase (end of ATB or water treatment) as described above. A total of 306 samples were collected to track individual changes in the fecal microbiome as a result of drug treatment. Samples were immediately frozen, transferred on dry ice to the Laboratory of Anaerobic Microbiology of the Institute of Animal Physiology and Genetics of the Czech Academy of Sciences (Czech Republic), and stored at -25 °C until use.

#### 2.4.2. DNA Extraction and PCR Amplification

Genomic DNA was isolated from fecal samples using the DNeasy PowerSoil Kit (Qiagen, Hilden, Germany) according to the manufacturer’s protocol. The concentration and purity of the extracted nucleic acids were checked using a NanoDrop 2000c UV-Vis spectrophotometer (Thermo Scientific, Waltham, MA, USA). DNA extracts were stored at -25 °C until use. The hypervariable regions V4-V5 of bacterial 16S rRNA were amplified with primers BactB-F: GGATTAGATACCCTGGTAGT and BactB-R: CACGACACGAGCTGACG according to Fliegerova et al. [79] using EliZyme HS Robust MIX Red (Elisabeth Pharmacon, Croydon, UK). Thermal cycling conditions were as follows: 5 min initial denaturation at 95°C; 30 cycles of denaturation at 95°C for 30 s, annealing at 57°C for 30 s and elongation at 72°C for 30 s; final elongation at 72°C for 5 min.

#### 2.4.3. High-throughput Sequencing

PCR amplicons (∼300 bp) were purified using the QIAquick PCR Purification Kit (Qiagen, Hilden, Germany) and libraries were prepared using the NEBNext Fast DNA Library Prep Set for Ion Torrent (New England BioLabs, UK) and the Ion Xpress Barcode Adapters 1-96 Kit (Thermo Fisher Scientific, Waltham, MA, USA). The amplicons were then pooled in equimolar ratios based on the concentration determined using a KAPA Library Quantification Kit (KAPA Biosystems). The sequencing template was prepared by emulsion PCR in a OneTouch 2 instrument using an Ion PGM OT2 HiQ View kit (ThermoFisher Scientific, Waltham, MA, USA). HTS was performed in an Ion Torrent PGM platform with an Ion 316 Chip Kit v2 BC (ThermoFisher Scientific) using an Ion PGM Hi-Q View Sequencing Kit (Thermo Fisher Scientific, Waltham, MA, USA) according to the manufacturer’s protocols.

#### 2.4.4. Bioinformatics and Statistical Analysis

Sequence data retrieved from the Ion Torrent Software Suite in fastq format were analyzed following the procedure of Bolyen et al. [80]. Reads were processed as described previously [81]. Briefly, sequences were trimmed, quality filtered, and chimeras were removed. The Amplicon Sequence Variants (ASVs) were generated using DADA2 software. To enable equal sampling depth, the dataset was subsampled to a minimum of 6000 reads per sample. Sequence processing resulted in the elimination of 10 samples, and the following 296 samples of respective groups of animals were included in the bacterial community structure analysis: the baseline LPS (n=76) and CTL (n=72) groups, test phase antibiotic-treated groups CTL/ATB (n=39) and LPS/ATB (n=43), and the water-treated groups CTL/W (n=33) and LPS/W (n=33). The VSEARCH-based consensus classifier was used to assign taxonomy against Greengenes database version 13_8 [82]. Bacterial community alpha diversity was assessed using Chao1, evenness, Faith’s phylogenetic diversity, and Shannon index. Beta diversity was assessed using Jaccard’s distance metric with Qiime2 version 2020.2 [80]. Sequence data and information were deposited in the Sequence Read Archive under accession number PRJNA765490.

The bacterial community was assessed using the parameters of alpha and beta diversity. Alpha diversities among groups of differently treated animals were compared using the Kruskal-Wallis H test. Statistical p-values and q-values with Benjamini-Hochberg false discovery rate are reported. Beta diversities among the studied groups were assessed with a nonparametric permutational multivariate ANOVA (PERMANOVA) test. The PERMDISP test was performed to support the PERMANOVA results. Linear discriminant analysis (LDA) effect size with standard parameters [83] was used to determine bacterial taxa with significantly different abundance levels.

## 3. Results

### 3.1. Body Weight

Body weight gain was significantly suppressed during neonatal LPS treatment (since PD 6) as shown by the results of ANOVA (PD 6: F(1,151)=63.57, p < 0.0001; PD 7: F(1,151)=77.19, p < 0.0001; PD 8: F(1,151)=94.37, p < 0.0001, PD 9: F(1,151)=94.01, p < 0.0001) (Fig. 2(A)). The significant negative effect of endotoxin persisted until weaning (PD 16-28) (PD 16: F(1,24)=10.56, p=0.0034; PD 21: F(1,24)=8.67, p=0.0071; PD 28: F(1,86) = 4.74, p=0.0315), but was diminished during further development of the animals (Fig. 2(B)). The significant changes in body weight were induced by the 10-day treatment with the ATB cocktail (PD 70: F(1,149)=6.19, p=0.0140) (Fig. 2(C)). A lower weight gain in both groups (CTL/ATB, LPS/ATB) compared to the respective controls (CTL/W, LPS/W) was clearly documented. However, a post-hoc analysis showed only a difference between the LPS/ATB and CTL/W groups (t=-3.28 df=75, p=0.0016). The body weight development in adulthood was not affected by early LPS administration (Fig. 2(C)), as the lower weight gain from weaning (PD 28) to adulthood (PD 70) of animals in the LPS/W group was not significantly different from the control group (CTL/W).

**Figure 2.**
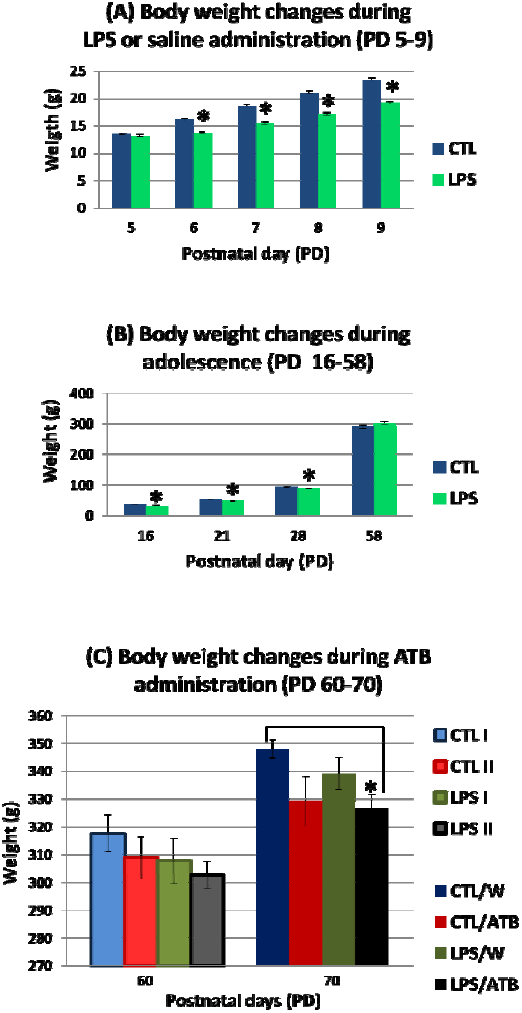
Body weight gain of rats during neonatal LPS treatment (A), during adolescence (B), and before (PD 60) and after (PD 70) 10-day ATB cocktail treatment (C). Results are expressed as mean ± SEM for the respective animal groups, *p □ 0.05.

### 3.2. Behavioral tests

All animals were subjected to behavioral tests in the following order: open field test, social recognition, and prepulse inhibition with respect to the ethological approach (i.e., all animal groups were tested in a mixed order; always with litter-matched controls). All animal groups were tested both in random order and during the same daytime period from 6:30 am to 2:00 pm, with one test per day.

#### 3.2.1. Open field test

In the open field pretest (i.e., before administration of the antibiotics), rats neonatally treated with the toxin showed significantly reduced locomotor activity (F(1, 44)=9.09; p=0.0043). The rats in the LPS I group traveled a shorter distance than the untreated CTL I group during 30 min, as shown in Figure 3(A) (pretest) and supported by post-hoc analysis (t=-3.48, df=21, p=0.0021). In the test session (i.e., at the end of antibiotic administration), no significant differences in distance traveled were observed between the 4 groups (CTL/W, CTL/ATB, LPS/W, and LPS/ATB) (Fig. 3(A), test), indicating that neither LPS (F(1,44)=1.38, p=0.2466) nor the ATB cocktail (F(1,44)=0.45, p=0.5056) affected the locomotor activities of the rats.

**Figure 3.**
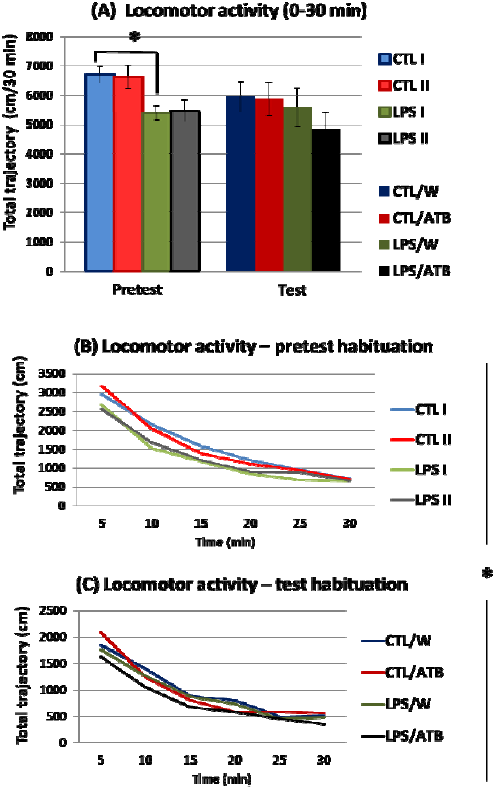
The spontaneous exploratory activity of differently treated groups of adult rats studied in the open field pretest (PD58/60) and test (PD68/70). The locomotor activity of the rats during the 30 min of the two sessions, expressed as total trajectory (A). Results are given as mean ± SEM for each animal group, *p □ 0.05. The habituation curves of the rats in the six 5-min blocks of the pretest (B) and test (C) sessions. Values are expressed as the mean distance traveled within each 5-min block.

On the other hand, the comparison of the habituation curves between the groups of animals showed a different locomotor activity of the rats treated with LPS (F(1,44)=6.84, p=0.0121) in the pretest and test sessions (i.e., mean values from 6 and 6 blocks of measurements taken during the pretest and test, respectively). The toxin-treated groups displayed a lower score of total distance traveled. Statistically significant differences in six 5-min measurement blocks (F(5,220)=2.53, p=0.0297) documented different dynamics of the habituation curve in the LPS-treated rats. Panels B and C in Figure 3 show that the curves of the LPS groups (LPS I and LPS II in pretest, LPS/W, LPS/ATB in test) are not parallel to the corresponding curves of the CTL groups (CTL I and CTL II in pretest, CTL/W and CTL/ATB in test), indicating faster habituation of the control groups. Thus, neonatal LPS treatment induced total hypoactivity in adult rats, especially in the open field pretest (Fig. 3(B)), whereas ATB administration had no significant effects on spontaneous exploratory activity and habituation parameters.

#### 3.2.2. Social recognition

Total exploration and social investigation of the juveniles were examined by two expositions during the pretest and test sessions (in total four juvenile object expositions). Both sessions resulted in similar social contact of all tested rat groups with the conspecific juvenile. Thus, social memory performance did not appear to be affected by both LPS administration in the early neonatal period and the 10-day ATB treatment in adulthood (data not shown).

However, the different behavior was observed in relation to rearing, which is used as an indicator of anxiety-like behavior. While no significant differences were observed in the pretest, the differences were evident in both expositions of the test (Fig. 4). Two-way ANOVA showed the influence of both treatments, i.e. LPS and ATB, indicating that LPS induced an increase in rearing, while the effect of ATB was opposite. This is already evident at the first exposure of the juvenile, where the LPS/W group had a significantly higher rearing frequency, as shown by the post hoc analysis of the difference between the CTL/ATB and LPS/W groups (t=-3.35, df=26, p=0.0025). At the re-exposition, after 30 min, no significant differences were found between the LPS/W and CTL/W groups. The ATB treatment alone (CTL/ATB) significantly decreased rearing frequency (F(1,26)=7.23, p=0.0123), while LPS insult in combination with ATB administration (LPS/ATB) significantly increased this behavior. This influence on rearing frequency was confirmed by post-hoc analysis, which showed significant differences between the CTL/ATB and CTL/W groups (t=3.30, df=26, p=0.0028) and the LPS/ATB and CTL/ATB groups (t=-2.58, df=26, p=0.0160).

**Figure 4.**
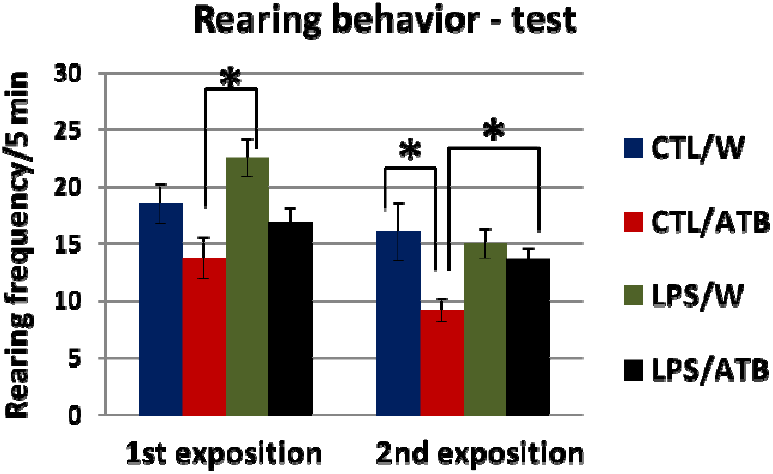
Rearing behavior of adult rats in social recognition test consisting of two expositions of the same juvenile subject. Results are expressed as mean ± SEM for each group of animals, *p □ 0.05.

#### 3.2.3. ASR and PPI

Acoustic startle responses did not differ significantly between animal groups in either session, indicating that neither early life LPS insult of rats by LPS nor antibiotic treatment of adults affected acoustic reactivity (Fig. 5(A)). Drug treatment resulted in only mild nonsignificant hyper-responsiveness to the startle pulse in animals in the LPS/ATB group (Fig. 5(A), test). However, comparison of pretest and test values revealed a significant difference between these two datasets showing higher acoustic sensitivity of all animal groups in the test session (pretest vs. test: F(1,57)=16.98, p<0.0001) and higher sensitivity to strong acoustic stimuli elicited by the ATB treatment (F(1,57)=6.93, p=0.0109) (CTL/ATB, LPS/ATB).

**Figure 5.**
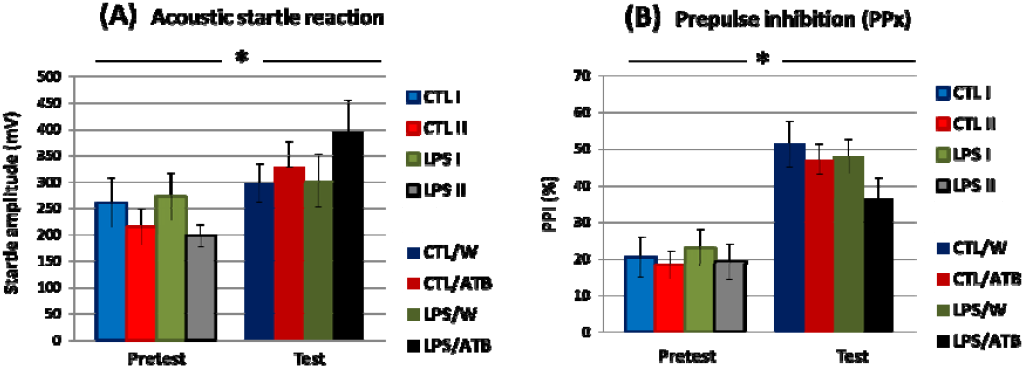
Acoustic startle response (A) and prepulse inhibition (B) of differentially treated groups of adult rats assessed in pretest (PD58/59) and test (PD68/69) sessions. PPx represents the mean of all different prepulses (PP3, PP5, and PP10). Results are given as mean ± SEM for each group of animals, *p □ 0. 05.

Prepulse inhibition was not significantly different between groups at both pretest and test, suggesting that neonatal LPS treatment and early adult ATB administration had no effect in adulthood. However, comparison of the pretest and test values revealed again a significant difference between these two datasets showing the increased reactivity of all groups in the test session (Fig. 5(B)) (pretest vs. test: F(1,57)=93.67, p<0.0001). Both tests indicate that the animals’ previous experience with the experimental setup resulted in a better response of the rats to the stimuli, regardless of the type of treatment. However, it appears that rats treated with LPS/ATB scored worse on PPI than animals of other groups.

### 3.3. Microbiome Analysis

In the present study, two hundred and ninety-six fecal samples were analyzed for bacterial diversity. Baseline samples collected from 148 animals at PD 60 (CTL=CTL I+CTL II, LPS=LPS I+LPS II) and test samples collected from 148 rats at PD 70 (CTL/W, LPS/W, CTL/ATB, and LPS/ATB) were analyzed for bacterial community composition to evaluate the effects of neonatal LPS insult and early adult ATB insult on the fecal microbiome in adulthood.

#### 3.3.1. Alpha and Beta Diversity

Bacterial community structure in feces from differentially treated rats was qualitatively and quantitatively examined for species richness and evenness, Faith’s phylogenetic diversity, and Shannon entropy. Alpha diversity metrics indicated a significant effect of treatment with the antibiotic cocktail, which resulted in a significant decrease in diversity in the CTL/ATB and LPS/ATB groups of animals, as shown in the alpha diversity boxplots (Supplementary Figure S1). Kruskal-Wallis pairwise tests for all calculated metrics showed significant differences (0.001 ≤ p c 0.05) between the two ATB-treated animal groups (CTL/ATB and LPS/ATB) and all other non-antibiotic groups (CTL, CTL/W, LPS, LPS/W). The decrease in diversity in the antibiotic-treated samples was similar in both ATB groups (CTL/ATB and LPS/ATB) and no significant difference was observed between them, indicating the crucial influence of the antibiotic cocktail regardless of the previous LPS treatment. No significant differences were found between the CTL and LPS groups (PD 60) and between the CTL/W and LPS/W groups (PD 70), indicating that the animals recovered from the toxin-induced insult at an early age (PD 5-9). Shannon entropy values determined by the pairwise Kruskal-Wallis H test are summarized for all rat groups in Table S1. Beta diversity, which assesses the similarity of bacterial communities between rat groups, was determined using Jaccard’s nonphylogenetic distance matrix. The main separation on axis 1 was driven by ATB treatment (Figure 6). The CTL/ATB (purple) and LPS/ATB (blue) groups clustered together (non-significant differences PERMANOVA p=0.081) and were significantly separated from four other rat groups (PERMANOVA p=0.001). Even if non-antibiotic groups clustered together, the community structures of samples collected at PD 60 were significantly different from those collected at PD 70, i.e., the CTL group (green) was different from the CTL/W group (yellow) (PERMANOVA p=0.001), and the LPS group (red) was different from the LPS/W group (orange) (PERMANOVA p=0.001). This difference can most likely be attributed to the sweetening of the water used to suppress the bitter taste of the antibiotics, which was applied to all animals, including the control groups. On the other hand, samples collected on the same day did not differ significantly, i.e., at PD 60, the control CTL group could not be distinguished from the LPS group (PERMANOVA p=0.596) and at PD 70, the control CTL/W group could not be distinguished from the LPS/W groups (PERMANOVA p=0.314). Pairwise PERMANOVA and PERMDISP results for all groups of rats are summarized in Table S1.

**Figure 6.**
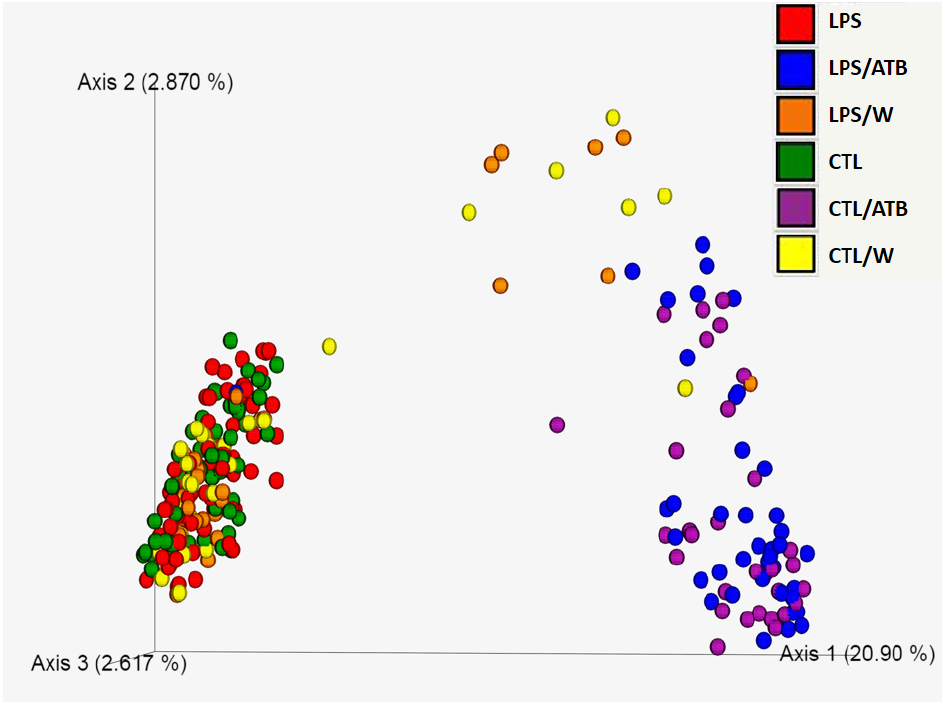
Principal coordinate analysis (PCoA) showing the Jaccard’s distance matrix of bacterial 16S rRNA amplicons from fecal samples from six groups of differently treated rats. Each dot represents one sample, and each rat group is indicated by a different color. The percentage of variation explained by the plotted principal coordinates is indicated on the axes.

### 3.2. Taxonomical Composition

A total of 9 phyla (including 110 bacterial phylotypes) were detected in the rat groups studied, but only 4 of them, including Firmicutes, Bacteroidetes, Proteobacteria, and Actinobacteria, had meaningful relative abundance (Fig. 7(A)). The abundances of the phyla Verrucomicrobia, Cyanobacteria, Deferribacteres, Tenericutes, and TM7 were considerably low (□ 0.03%) and are summarized as “others” in Figure 7(A). Firmicutes was detected as the dominant phylum in all groups (45.6-57.9%) regardless of treatment. Bacteroidetes was the second dominant phylum in the non-antibiotic-treated groups (19.7-31.2%), followed by a low abundance of Actinobacteria (0.3-1.8%), as shown in Figure 7(A). In both ATB-treated groups (CTL/ATB, LPS/ATB), Proteobacteria (52%), dominated by order Enterobacteriales (25.1-30.1%) and Burkholderiales (13.0-14.4%), were the second most abundant phylum (40.4-43.2%), while Actinobacteria were depleted (Figure 7(A, B, C)). In the non-antibiotic groups, Firmicutes and Bacteroidetes were mainly represented by the order Clostridiales (30.6-48.3%) and Bacteroidales (19.7-31.2%), respectively, and a high relative abundance of unassigned bacteria (11.4-20.4%) was observed in all four groups (CTL, LPS, CTL/W, LPS/W). The ATB treatment completely suppressed Clostridiales, while Lactobacillales were induced and formed the most abundant order (51.4-53.2%), as shown in Figure 7(C). The reduced bacterial diversity due to antibiotic treatment of animals (CTL/ATB, LPS/ATB) is well seen at the family level (Figure 7(D)). Both ATB-treated animal groups were dominated by *Lactobacillaceae* (51.4-53.2%) represented by the genus *Lactobacillus, Enterobacteriaceae* (26-31.1%) represented by *Proteus* and unassigned genus, and *Alcaligenaceae* (13-14.4%) represented by the genus *Sutterella*. ATB treatment considerably suppressed the abundance of unassigned bacteria (0.5-0.7%).

**Figure 7.**
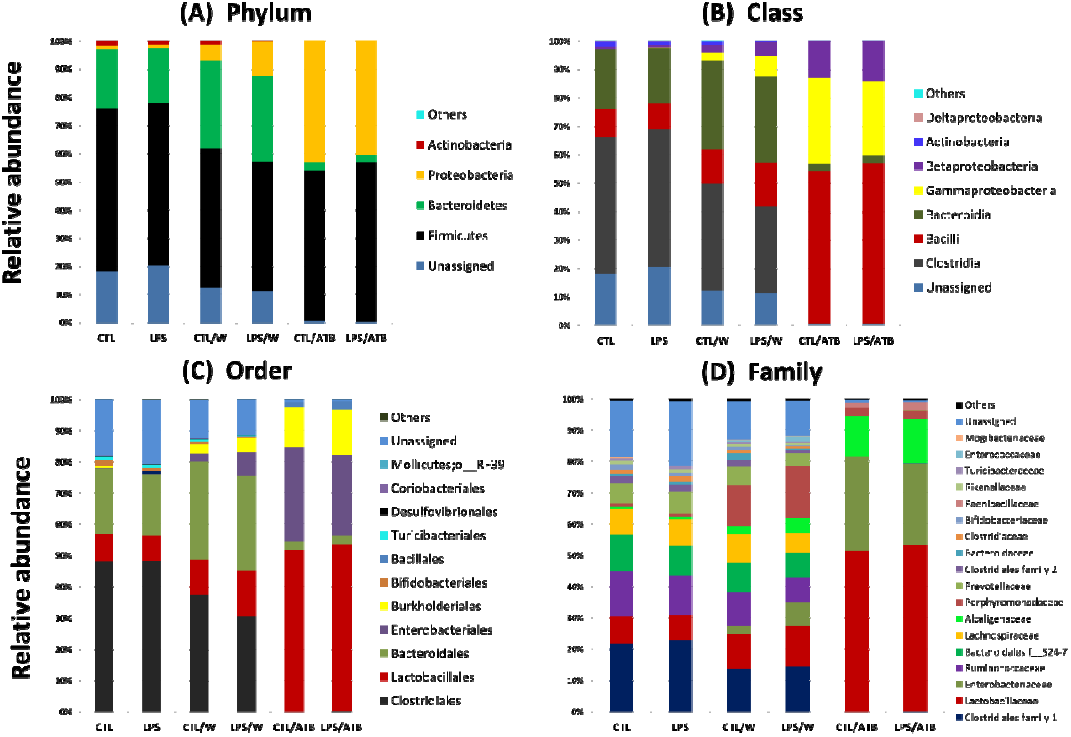
Relative abundance of fecal bacteria at the phylum (A), class (B), order (C), and family (D) levels in six groups of differently treated rats. Only taxa with a relative abundance of ≥ 0.1 % in at least one group are reported.

To elucidate specific phylotypes that respond to LPS and/or ATB treatment, the linear discriminant analysis (LDA) effect size (LEfSe) algorithm was applied to identify taxa that are more abundant in one group than in the other corresponding group of animals. To identify specific phylotypes that respond to LPS and/or ATB treatment, the linear discriminant analysis (LDA) algorithm was applied to determine the effect size (LEfSe) to identify taxa that are more abundant in one group than in the other group of animals. Regarding the influence of LPS, the four bacterial taxa were differentially abundant in the CTL/W and LPS/W groups (LDA score > 3.0). *Oscilospirra* was overrepresented in the LPS/W group, whereas *Bacteroides*, Actinomycetales, and *Coriobacteriaceae* were overrepresented in the control CTL/W group, as shown by a cladogram (Fig. 8(A)) and a histogram (Fig. S2).

**Figure 8.**
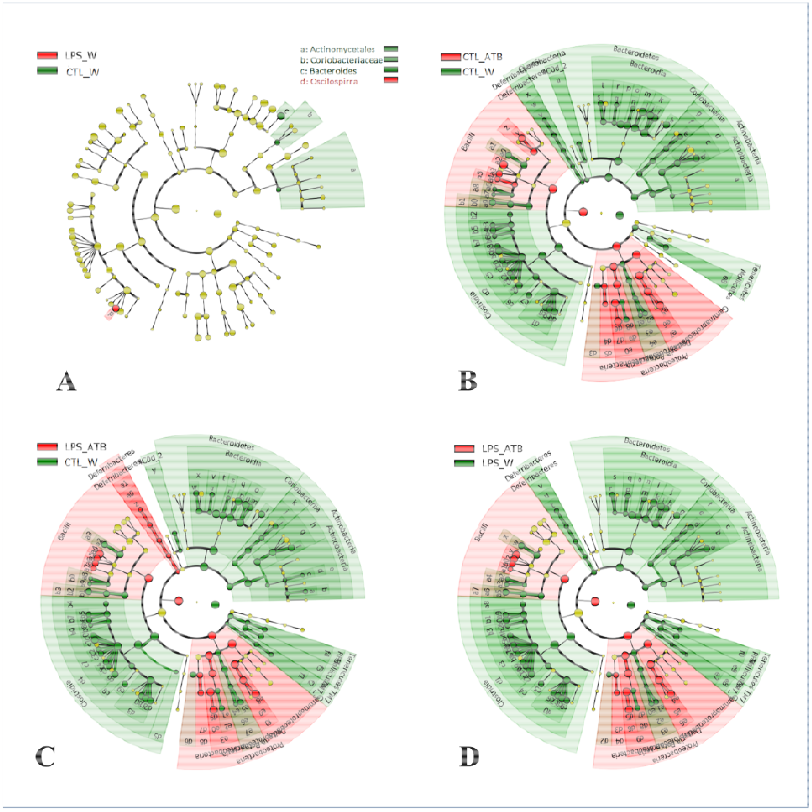
Cladograms generated with LEfSe software show taxa with different abundance in fecal samples from rats treated with LPS and/or an antibiotic cocktail compared with their respective controls. Regions in red indicate taxa that were enriched in LPS- and/or antibiotic-treated animals, whereas regions in green indicate taxa that were enriched in the respective control groups. (A) Four differentially abundant bacterial phylotypes between the groups of LPS-treated rats and the control group (LPS/W vs. CTL/W). (B) Ninety-five differentially abundant bacterial phylotypes between the group of antibiotic-treated rats and the control group (CTL/ATB vs. CTL/W). (C) One hundred and three differentially abundant bacterial phylotypes between the group of rats treated with LPS and antibiotics and the control group without any treatment (LPS/ATB vs. CTL/W). (D) Ninety-eight differentially abundant bacterial phylotypes between the group of rats treated with LPS and antibiotics and the control group treated with LPS (LPS/ATB vs. LPS/W). The names of the significantly enriched phylotypes are indicated in the LEfSe histograms of the LDA scores (Supplementary Figures S2-S4).

However, no significant differences were found between the LPS and CTL groups in PD 60 (data not shown). Regarding the influence of the antibiotic cocktail, the 95 bacterial taxa differed significantly in abundance between the CTL/ATB and CTL/W groups (LDA score > 2.4), as shown in Fig. 8(B) and Fig. S3. ATB treatment (CTL/ATB) enriched 19 phylotypes of the phylum Proteobacteria and the class Bacilli. Among the 76 taxa enriched in the control group (CTL/W), the phylotypes of Bacteroidetes, Clostridia, and Actinobacteria are meaningful, whereas the taxa of Tenericutes, Defferibacteria, and Cyanobacteria are negligible due to their very low relative abundance (together □ 0.03%). The combined treatment with LPS and ATB resulted in 103 and 98 differentially expressed phylotypes, respectively (LDA score > 2.4), based on the comparison of the LPS/ATB group with the CTL/W (Fig. 8(C) and Fig. S4(A)) and LPS/W (Fig. 8(D) and Fig. S4(B)) groups, respectively. The overrepresented phylotypes are similar to those of the antibiotic-only treated group (CTL/ATB). Thus, the observed effect can be mainly attributed to the application of the antibiotic, which is confirmed by the limited changes in the LPS/W group (Fig. 8(A)) and the nonsignificant changes in LPS-induced taxa abundance determined by LEfSe analysis of the LPS and CTL groups (PD 60) and the CTL/ATB and LPS/ATB groups (PD 70).

## 4. Discussion

Here, we use an animal model to try to determine whether the LPS challenge in early life, followed by an ATB insult in early adulthood, can induce the behavioral changes relevant to schizophrenia. However, the general results of our study suggest that this combination of two stressors did not have significant effects on the behavior of the rats. The strong tendency toward higher vulnerability to acoustic reflex and lower levels of PPI in the LPS/ATB-treated group was evident but not statistically proven. This finding is not consistent with the schizophrenia-like models of several authors [17,32,34–36,84], all of whom described an LPS-induced deficit in prepulse inhibition in adult animals. However, in contrast to our work, these studies used either neonatal intrahippocampal [32] or prenatal intraperitoneal LPS injections [17,34–36] suggesting that the injection site and timing of LPS application appear to be critical for subsequent behavioral changes. Surprisingly, there is no information on the effect of antibiotic treatment and the resulting disruption of the microbiome on sensorimotor gating, as animals in the relevant (antibiotic-using) studies [51,53,85,86] have not been tested for prepulse inhibition and/or responses to intense acoustic stimuli. The usefulness of including such tests is illustrated by the work of Giovanoli et al. [44], who show that antibiotic treatment can prevent the development of sensorimotor gating deficiency induced by combined prenatal immune activation and peripubertal stress in a mouse model of schizophrenia. This finding may suggest that the antibiotics used as a second stressor in our study may have acted in the opposite direction and protected the animals from the consequences of LPS-induced inflammation. However, the results of the LPS-treated group (without ATB treatment) suggest that the effects of the early life immune insult was transient and adult rats generally overcame the endotoxin-induced discomfort documented in young rats by significantly reduced weight gain by weaning. No downstream effects of LPS on social behavior and sensorimotor gating in rats were detected in their adulthood. On the other hand, LPS-challenged rats exhibited locomotor hypoactivity, but only in the pretest, lower habituation ability in OFT, and increased frequency of rearing as an indicator of anxiety-like behavior in the presence of conspecific young. These results suggest some long-lasting consequences of LPS treatment, but the effect of endotoxin insult was not as complex as in other studies. While the suppressive effect of LPS on weight gain [24,78,87–91] and locomotor activity [24,27,88,92–98] is well known, the depressive effect of LPS on social exploration of rats described by many authors [24,87,88,92] was not observed in our study.

No negative changes were observed in the fecal bacterial population of LPS-treated animals, again suggesting that adult rats overcame the early age endotoxin insult. Increased occurrence of *Oscillospira* (LPS/W, PD 70) may even be considered beneficial because this bacterium is known to produce butyrate [99], which is a major source of energy for colon epithelial cells, plays an important role in maintaining the stability of the gut microbiota, alleviates mucosal inflammation, and strengthens the epithelial defense barrier [100,101]. This bacterium commonly found in 60% of humans [102], is significantly decreased in patients with inflammatory bowel disease (IBD) [103,104] and was significantly reduced in SI rats (social isolation of rat pups from weaning as an animal model of schizophrenia) [105]. On the other hand, Xia et al. [61] described a higher abundance of this genus in adult LPS-treated mice immediately after 5 days of LPS challenge, which is not consistent with IBD studies in humans and seems to be in contradiction with the proinflammatory nature of LPS [106]. However, little is known about *Oscillospira* metabolism and/or physiology due to its recent discovery and difficult cultivation [99].

On the other hand, the effect of ATB on the microbiome was immense. A 10-day antibiotic treatment drastically reduced microbial diversity and altered the bacterial composition of adult rat feces. Both groups of ATB-treated animals were clearly separated from the non-antibiotic groups in PCoA analysis, regardless of LPS treatment. A significant decrease in bacterial diversity due to ATB treatment is consistent with the results of several studies in rats [53,107– 109], mice [51,54,110], and hamsters [46]. However, the disturbances of the microbial community in the aforementioned studies were highly dependent on the type of antibiotics administered and the study design. In our work, the most obvious changes were a significant increase in Proteobacteria, a decrease in Bacteroidetes, and a depletion of Actinobacteria, phyla with low abundance, and unassigned species. The relative abundance of Firmicutes was not altered, but the community composition within this phylum was notably affected. Proteobacteria is a common phylum of mammal GIT, ranging from 2.5 to 10% in the healthy rat intestine [111–113]. Its elevated abundance is however increasingly recognized as a possible microbial signature of the disease state [114]. In our study, Proteobacteria were represented at the lowest taxonomically identifiable level by the family *Enterobacteriaceae* and the genera *Proteus* and *Sutterella*. The family *Enterobacteriaceae* includes proinflammatory, potentially pathogenic bacteria [115], *Proteus* species possess many virulence factors potentially relevant to gastrointestinal pathogenicity [116,117], and *Sutterella* was found to be associated with GI infections [118] and inflammatory bowel disease [119–121]. The increased abundance of *Sutterella* spp. has also been detected in the feces of children with autism spectrum disorders (ASD) [122–124], indicating the possible role of this bacterium in neurodevelopmental diseases. The effect of antibiotics on the Firmicutes phylum resulted in extensive suppression of the diverse bacterial diversity (found in the groups without antibiotics) and monoculture of the genus, which accounted for 94% of the Firmicutes in the LPS/ATB group. Antibiotics also depleted the majority of Clostridia and eliminated the *Ruminococcaceae* and *Lachnospiraceae* families, which are generally considered beneficial for gut health [125]. These effects together with the diminishing of Bacteroidetedes strongly suggest gastrointestinal disturbances induced by ATB treatment. Therefore, it was surprising that the observed suppression of bacterial diversity had no effect on rat behavior. These results are in contradiction with data from the rodent literature showing the negative effect of ATB on spatial recognition memory in rats [53,109], ATB-induced cognitive deficits [110], disruption of novel object recognition memory [54], increased exploratory behavior [126], increased immobility and reduced social recognition [127] in mice, and a reduction in the duration of nose-to-nose investigation after antibiotic treatment in hamsters [46]. On the other hand, Hoban et al. [53] observed that antibiotic treatment of adult rats had no effect on anxiety-related behavior assessed using Elevated Plus Maze (EPM) and Open Field (OF) tests. Cross-comparison of experimental results is difficult and sometimes problematic because results are strongly influenced by many variable factors, such as type and duration of drug treatment, type of drug administration and dose, age of animals at time of insult/test, type of behavioral tests, breed of rodents, number of animals in an experiment, source of animals, and even factors related to rodent husbandry. Our study design is similar to that of the hamster study by Sylvia et al. [46]. Antibiotic treatment of adult hamsters strongly affected gut microbial communities, but this effect was not influenced by LPS insult at early age, similar to our study. On the other hand, LPS-treated male hamsters exhibited anxiety-like behaviors after antibiotic treatment in adulthood, a phenomenon that was not observed in our experiments.

In conclusion, the two-hit model tested in this study did not induce schizophrenia-like behavior in adult male rats. Apart from reduced weight gain, there were no behavioral abnormalities caused by neonatal LPS immune challenge combined with an early adult antibiotic insult. A sensorimotor gating deficit, considered a behavioral phenomenon relevant to schizophrenia, was observed, but the difference between the two-hit-treated group and the control group was not statistically significant. Our results are quite surprising in view of the well-documented negative influence of early age endotoxin treatment and the reported negative effects of gut microbial disruption on rodent behavior. LPS-only treated animals showed in adulthood hypoactivity and induced anxiety-like behavior in the social recognition paradigm, but these behavioral changes were not exacerbated by ATB-induced gut dysbiosis. In animals treated with ATB alone, the gut bacterial community was seriously disrupted, but the dysbiosis did not affect behavioral responses in the tests. The protective, antiinflammatory effect of ATB administration cannot be supposed due to the significant increase in potentially harmful bacteria and elimination of beneficial microbes. The lack of interaction between LPS and ATB treatment suggests that there may be independent mechanisms of action for both insults. Comparison with data from the literature suggests that targeting and timing of hits have a large impact on outcomes. Further work is needed to better understand how an immune challenge and microbiome dysbiosis may influence animal health status and behavior.

## Supplementary Materials

Table S1: Shannon diversity values evaluated by pairwise Kruskal–Wallis *H* test and pairwise PERMANOVA and PERMDISP results based on the Jaccard distance matrices for all groups of rats. Figure S1: Alpha diversity boxplots represented by Shannon entropy (A), observed ASVs (B), and Faith’s phylogenetic diversity (C) for differently treated groups of rats showing significantly decreased metrics for groups of animals treated by antibiotic cocktail (LPS/ATB, CTL/ATB). Figure S2: Linear discriminant analysis (LDA) scores for 4 bacterial phylotypes with significantly different abundance in fecal samples between group of LPS treated rats (LPS/W) and control group (CTL/W). Figure S3: Linear discriminant analysis (LDA) scores for 95 bacterial phylotypes with significantly different abundance in fecal samples between group of antibiotic-treated rats (CTL/ATB) and control group (CTL/W). Figure S4: (A) Linear discriminant analysis (LDA) scores for 103 bacterial phylotypes with significantly different abundance in fecal samples between LPS and antibiotic-treated rats’
s group (LPS/ATB) and control group (CTL/W). (B) Linear discriminant analysis (LDA) scores for 98 bacterial phylotypes with significantly different abundance in fecal samples between LPS and antibiotic-treated rats group (LPS/ATB) and LPS treated control group (LPS/W).

## Author Contributions

Conceptualization, H.T., and K.F.; methodology, H.T., and K.F.; formal analysis and investigation, S.K., H.S., J.M., and J.K.; resources, H.T., K.F., J.K., and T.P.; data curation, S.K.; writing—original draft preparation, K.F., H.T., H.S., and J.K.; writing—review and editing, K.F., H.T., J.K., L.J., and T.P.; supervision, H.T., K.F., and S.K.; funding acquisition, T.P., H.T., J.M., and K.F. All authors have read and agreed to the published version of the manuscript.

## Data Availability Statement

The data presented in this study are openly available in the Sequence Read Archive under the accession number PRJNA765490.

## Acknowledgments

We would like to thank Dagmar Schierová for helpful advice during the HTS data analyses.

