## Supplemental material for "Antibiotic-induced dysbiosis does not potentiate the effect of lipopolysaccharide insult in male Wistar rats"

Table S1. Shannon diversity values evaluated by pairwise Kruskal–Wallis *H* test and pairwise PERMANOVA and PERMDISP results based on the Jaccard distance matrices for all groups of rats. Significant p-values are in red.

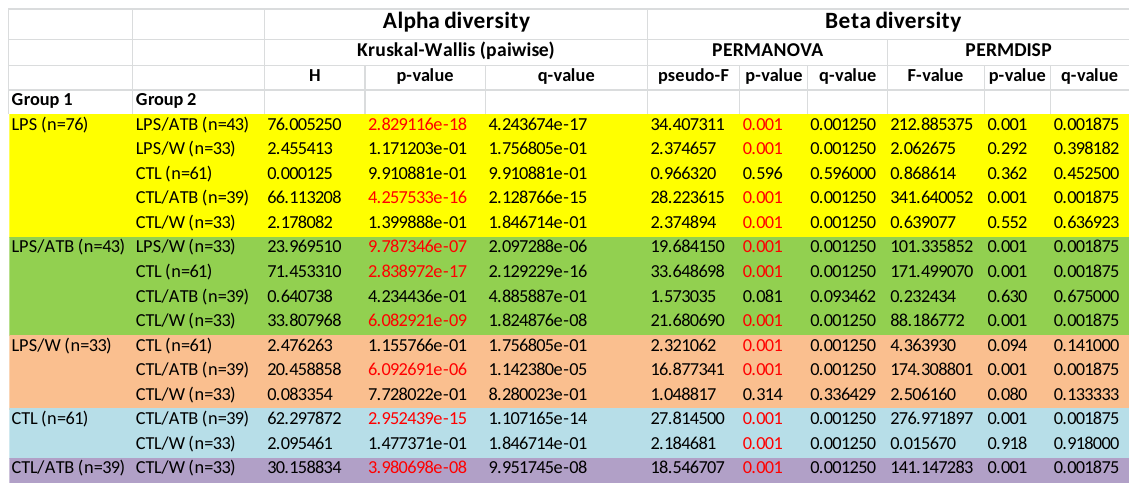

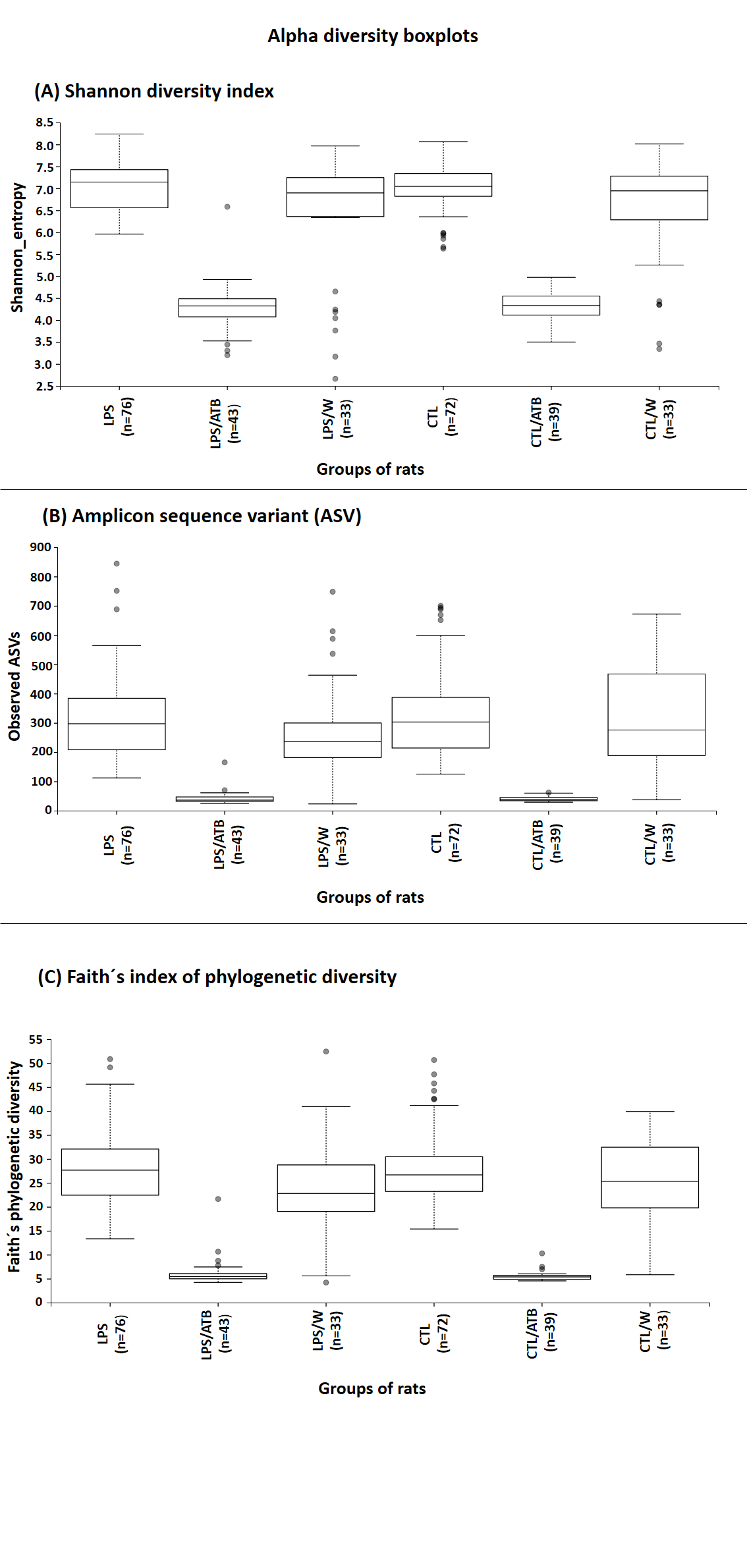

Figure S1. Alpha diversity boxplots represented by Shannon entropy (A), observed ASVs (B) and Faith's phylogenetic diversity (C) for differently treated groups of rats showing significantly decreased metrics for groups of animals treated by antibiotic cocktail (LPS/ATB, CTL/ATB). Number of rats inside each group is given in brackets.

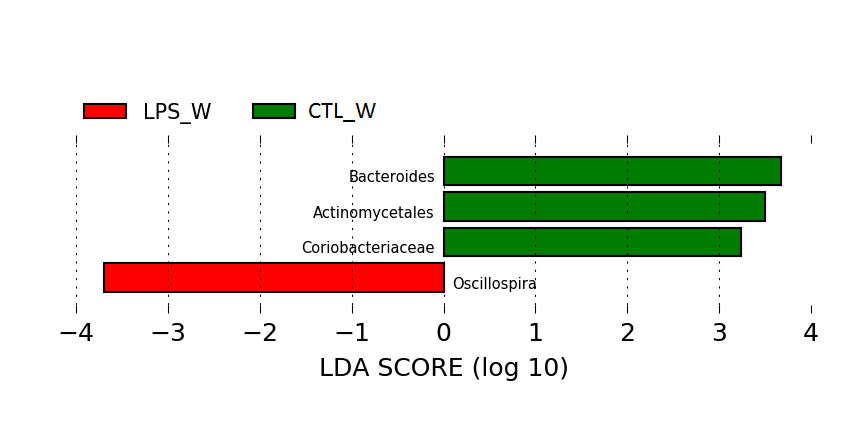

Figure S2. Linear discriminant analysis (LDA) scores for 4 bacterial phylotypes with significantly different abundance in fecal samples between group of LPS treated rats (LPS/W) and control group (CTL/W). The length of the bar represents the log10 transformed LDA score, indicated by vertical dotted lines. Negative (red bars) LDA scores represent bacterial taxa over-abundant in LPS/W group, positive (green bars) represent bacterial taxa over-abundant in CTL/W group.

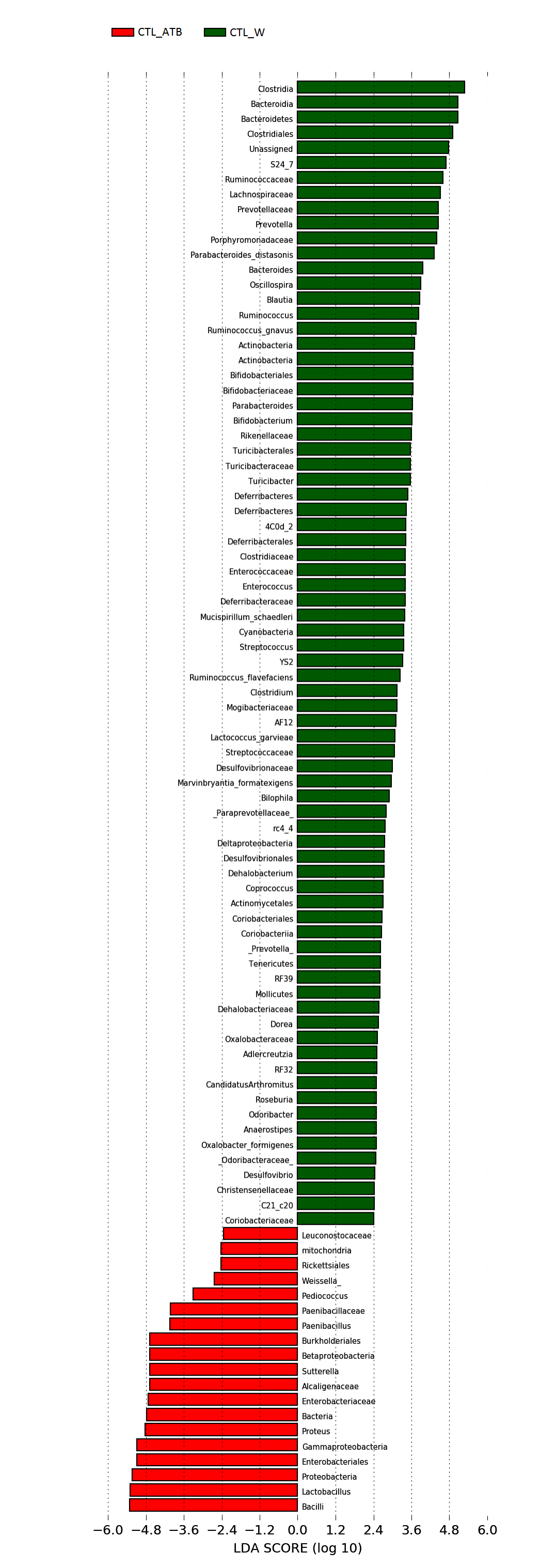

Figure S3. Linear discriminant analysis (LDA) scores for 95 bacterial phylotypes with significantly different abundance in fecal samples between group of antibiotic-treated rats (CTL/ATB) and control group (CTL/W). The length of the bar represents the log10 transformed LDA score, indicated by vertical dotted lines. Negative (red bars) LDA scores represent bacterial taxa over-abundant in CTL/ATB group, positive (green bars) represent bacterial taxa over-abundant in CTL/W group.

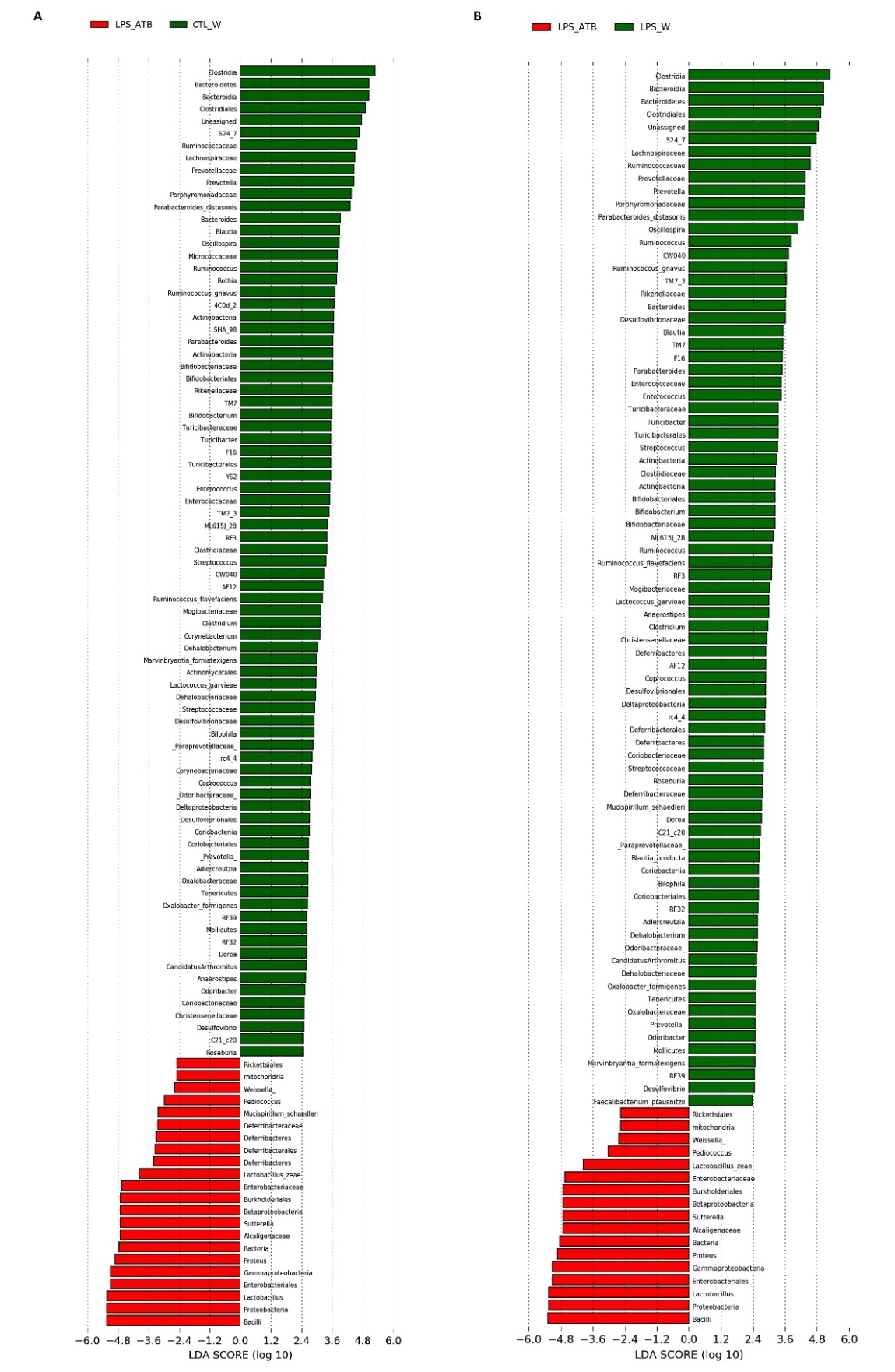

Figure S4. A. Linear discriminant analysis (LDA) scores for 103 bacterial phylotypes with significantly different abundance in fecal samples between LPS and antibiotic treated rats´ group (LPS/ATB) and control group (CTL/W). B. Linear discriminant analysis (LDA) scores for 98 bacterial phylotypes with significantly different abundance in fecal samples between LPS and antibiotic treated rats group (LPS/ATB) and LPS treated control group (LPS/W).The length of the bar represents the log10 transformed LDA score, indicated by vertical dotted lines. Negative (red bars) LDA scores represent bacterial taxas over-abundant in LPS/ATB group, positive (green bars) represent bacterial taxa over-abundant in CTL/W (A) and LPS/W group (B).
